# Hypermorphic *SERK1* mutations function via a *SOBIR1* pathway to activate floral abscission signaling

**DOI:** 10.1101/453944

**Authors:** Isaiah Taylor, John Baer, Ryan Calcutt, John C. Walker

**Author notes:** Present address: Department of Biology and Howard Hughes Medical Institute, Duke University, Durham, NC 27708, USA. Present address: Department of Medicine, Washington University in St. Louis, St. Louis, MO 63130, USA. Present address: Biology Department, Washington University in St. Louis, St. Louis, MO 63130, USA.

## Abstract

In Arabidopsis, the abscission of floral organs is regulated by two related receptor-like protein kinases (RLKs), HAESA and HAESA-like 2 (HAE/HSL2). HAE/HSL2, in complex with members of the SERK family of coreceptor protein kinases, are activated by the binding of the proteolytically processed peptide ligand IDA. This leads to expression of genes encoding secreted cell wall remodeling and hydrolase enzymes. *hae hsl2* mutants fail to induce expression of these genes and retain floral organs indefinitely. In this paper we report identification of an allelic series of *hae hsl2* suppressor mutations in the *SERK1* coreceptor protein kinase gene. Genetic and transcriptomic evidence indicates these alleles represent a novel class of gain of function mutations that activate signaling independent of HAE/HSL2. We show that the suppression effect surprisingly does not rely on protein kinase activity of SERK1, and that activation of signaling relies on the RLK gene *SOBIR1*. The effect of these mutations can be mimicked by loss of function of *BIR1,* a known negative regulator of *SERK*-*SOBIR1* signaling. These results suggest *BIR1* functions to negatively regulate *SERK-SOBIR1* signaling during abscission, and that the identified *SERK1* mutations likely interfere with this negative regulation.

## Introduction

Abscission is the process by which plants shed structures, such as fruit, leaves, and floral organs. Abscission occurs as the result of a developmental process or is triggered by damage or adverse environmental conditions. In Arabidopsis, abscission of sepals, petals, and stamen follows pollination in a developmentally programmed manner, while abscission of cauline leaves occurs as a result of drought stress or pathogen infection (1–4). Abscission is regulated by the two redundant leucine-rich repeat-receptor like protein kinases (LRR-RLKs) HAESA and HAESA-like 2 (HAE/HSL2) (5–7). Binding of the proteolytically processed, secreted peptide INFLORESCENCE-DEFICIENT IN ABSCISSION (IDA) induces association of HAE/HSL2 and members of the SOMATIC EMBRYOGENESIS RECEPTOR-LIKE KINASE (SERK) family of LRR-RLK co-receptors (8–10). This association activates a downstream MAP kinase cascade comprised of MAP kinase kinase 4 and 5 (MKK4/5) and MAP kinase 3 and 6 (MPK3/6) (6,9). This MAP kinase cascade targets transcriptional repressors of the AGAMOUS-like family, including AGL15, leading to de-repression of *HAE* expression and an increase in signaling via positive feedback (11–14). This signaling pathway regulates expression of a number of genes involved in cell wall remodeling, hydrolysis of cell wall and middle lamella polymers such as pectin (15). Plants with mutations in *HAE/HSL2* or *IDA,* or in high-order mutants of *SERK* genes, display floral abscission defects indicative of a failure to properly break down the middle lamella between the abscising organ and the body of the plant (6,9,16). Additionally, mutants in the ADP-Ribosylation Factor GTPase Activating Protein (ARF-GAP) gene *NEVERSHED* are also abscission deficient and possess a disorganized secretory system in the abscission zone (2). The abscission deficient phenotype of *nev* can be suppressed by mutations in *SERK1*, as well as mutations in the RLK gene *SUPPRESSOR OF BIR1-1/SOBIR1* (also known as *EVERSHED/EVR*), and the cytosolic RLK *CAST AWAY* (17,18). The exact cause of the *nev* mutant phenotype, as well as the mechanism of *nev* suppression, are not fully understood. Recent work has demonstrated *nev* mutants have ectopic lignification patterns in the abscission zone and exhibit widespread transcriptional reprogramming, including strong induction of biotic stress response gene expression (19,20). These results together suggest the *nev* phenotype may be related to mis-regulation of molecular signaling, possibly involving a pathway regulating lignification of the abscission zone.

In this work, we sought to understand additional regulators of abscission signaling using a genetic suppression approach. We identified an allelic series of putative gain of function *SERK1* mutations capable of suppressing the abscission defect of *hae hsl2.* These mutant alleles represent a novel class of hypermorphic *SERK* gene mutations that appear to signal through the RLK *SOBIR1,* likely by interfering with the function of the negative regulator of signaling *BIR1*. These alleles provide insights into the regulation of downstream signaling processes by SERK proteins. Implications for understanding of the *nev* mutant are discussed.

## Results

### Identification of *hae-3 hsl2-3* suppressor mutations in the *SERK1* gene

To identify regulators of abscission signaling, we performed suppressor screens of the previously described *hae-3 hsl2-3* abscission deficient mutant (15,21). We isolated three strong suppressors, an intermediate suppressor, and a weak suppressor. Initial mapping by sequencing of a strong suppressor identified a semi-dominant, linked missense mutation in the *SERK1* gene, substituting L326 for phenylalanine [Fig. 1A and Fig. 1B; Supp. Table S1]. Additional genetic analysis demonstrated all 5 mutants contained semi-dominant, linked missense mutations affecting conserved SERK protein residues [Fig. 1C; Supp. Table S1; Supp. Fig. S2].

**Figure 1:**
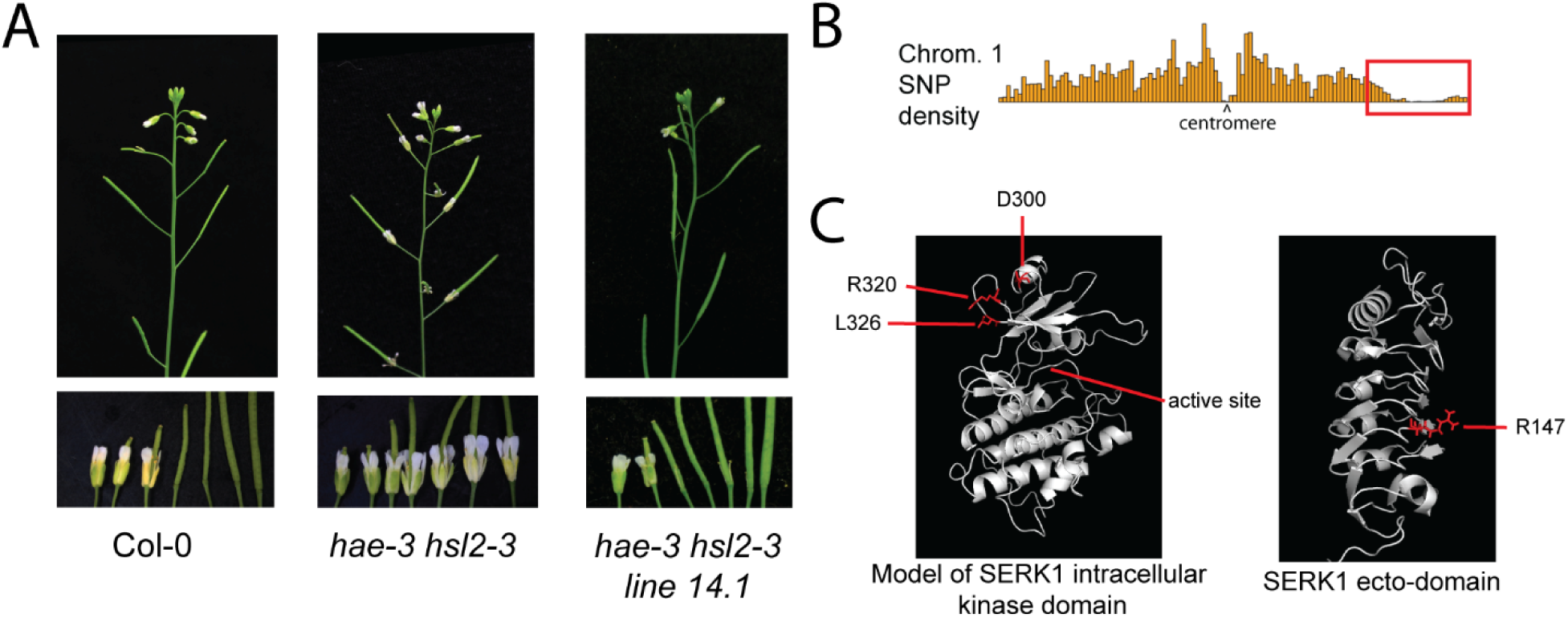
Phenotype and molecular mapping of *hae-3 hsl2-3* suppressor mutations. A) Floral abscission phenotype of Col-0, *hae-3 hsl2-3*, and *hae-3 hsl2-3* line 14.1 B) Landsberg erecta SNP density resulting from mapping by sequencing of bulked DNA from strongly abscising *hae hsl2* line 14.1 *(Col-0) x hae hsl2 (Ler)* F2 plants, indicating linkage to chromosome 1 C) Predicted location of *hae hsl2* suppressor mutations on SERK1 protein

To gain insight into the possible functions of the mutated residues, we mapped them onto a model of the SERK1 intracellular domain based on the crystal structure of the closely related BAK1/SERK3 protein kinase domain, as well as on the recently solved crystal structure of the SERK1 ecto-domain (22,23) [Protein Data Bank IDs: 3ULZ and 4LSC]. The four protein kinase domain mutations are localized to a patch of three surface exposed residues on the N-lobe of the protein kinase domain distal to the catalytic cleft [Fig.1C]. The phenotypes of these mutants are all strong or intermediate [Supp. Fig. S1; Supp. Fig. S2]. The extracellular domain mutation with the weak suppression phenotype maps to the ligand-receptor binding face of the ecto-domain [Fig. 1C]. This residue, R147, makes side chain interactions with the residue D123, which is homologous to BAK1 residue D122. Mutation of D122 yields a hypermorphic variant of BAK1 displaying ligand independent association with the brassinosteroid receptor BRI1, and reduced affinity for the ecto-domain of members of the BIR family of negative regulators of signaling (24,25). However, we found the weak phenotype of this mutant difficult to study, and instead focused on the intracellular domain mutants. Since the four strongest mutations cluster to a spatially restricted region of the protein kinase domain, we hypothesize that the mutations have a similar effect. To study this effect, we examined the *serk1-L326F* mutant as a representative allele.

### Genetic analysis of *hae-3 hsl2-3 serk1-L326F*

Because *SERK1* positively regulates abscission and because these mutations are all semi-dominant, we hypothesized that they are gain of function mutations. We performed a number of genetic experiments to test this hypothesis. We first crossed the *hae-3 hsl2-3* mutant with the well-characterized *serk1-1* loss of function T-DNA insertion mutant. Consistent with previously published analysis of *serk1* loss of function mutations, we found the *serk1-1* allele is unable to suppress the abscission defect of *hae-3 hsl2-3,* demonstrating specificity of the suppression defect to the semi-dominant alleles isolated in our screen (26) [Fig. 2A]. We next crossed the *serk1-L326F* mutant with the *hae-1 hsl2-4* double mutant. This mutant has a T-DNA insertion in the first exon of *HAE* and a premature stop codon in the first exon of *HSL2* and is a predicted protein null (21). We isolated the *hae-1 hsl2-4 serk1-L326F* triple mutant and found it displays near complete suppression of the abscission defect similar to *hae-3 hsl2-3 serk1-L326F,* demonstrating non-specificity of the suppression effect in regard to alleles of *hae* and *hsl2* [Fig. 2B]. We further performed a transgenic recapitulation experiment, where we transformed *SERK1pr::SERK1* wildtype and *SERK1pr::SERK1-L326F* mutant transgenes into *hae-3 hsl2-3*. In the T1 generation, we found that 0/16 plants transformed with the wildtype transgene displayed abscission [Fig. 2C; Supp. Table S3 for all transgene counts]. For the *SERK1pr::SERK1-L326F* transgene, 10/30 T1 plants displayed weak or strong suppression [Fig. 2C, Supp. Table S3]. These results are consistent with a model wherein *serk1-L326F* is a dose-dependent, hypermorphic mutation.

**Figure 2:**
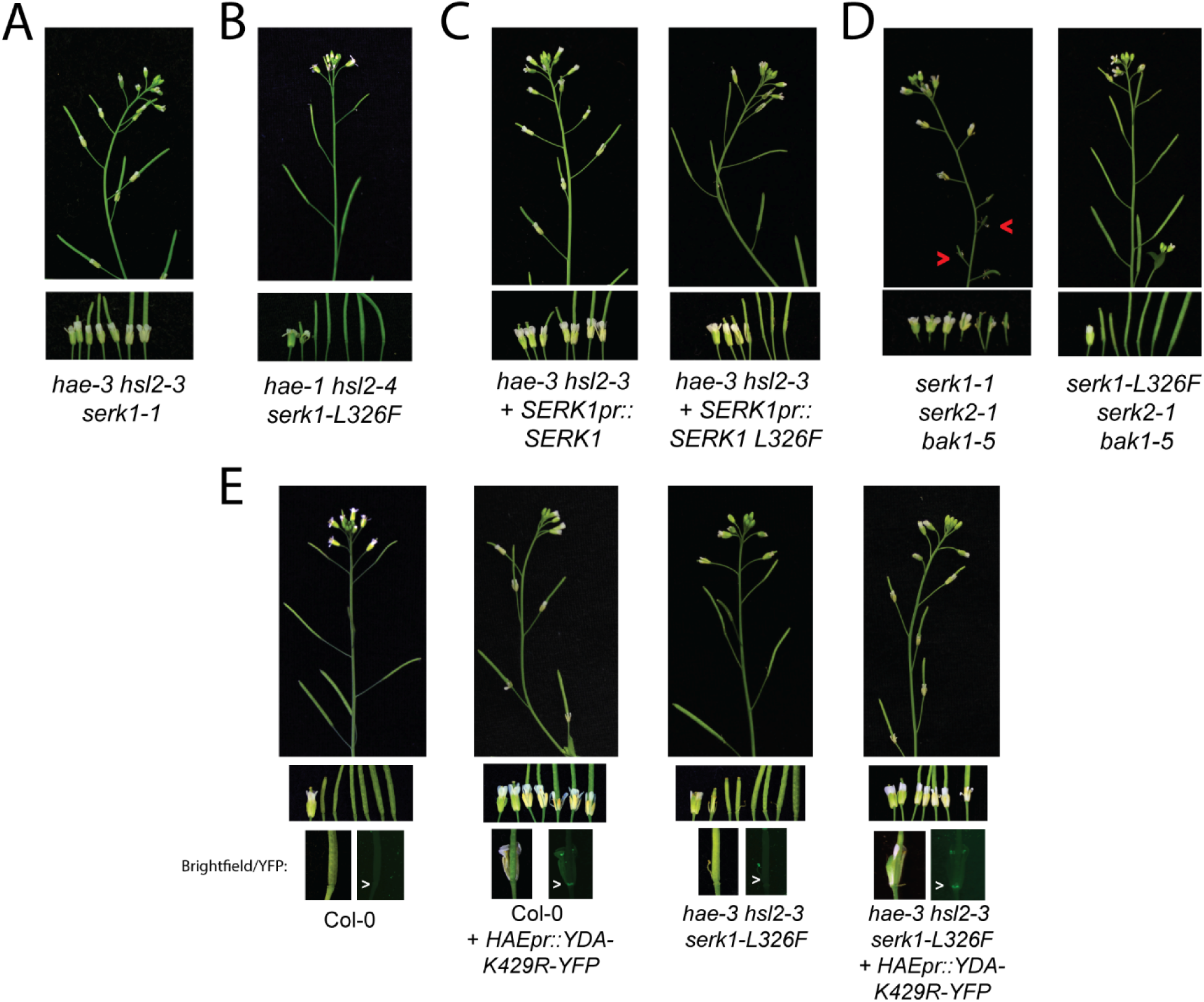
Genetic analysis of representative hypermorphic *SERK1* allele. A) Phenotype of *hae-3 hsl2-3 serk1-1* triple mutant B) Phenotype of *hae-1 hsl2-4 serk1-L326F* triple mutant C) Transgenic recapitulation of the *hae hsl2* suppression phenotype D) Phenotype of *serk1 serk2 bak1* triple mutants E) Phenotype of *HAEpr::YDA-YFP K429R* transformed into Col-0 and *hae-3 hsl2-3 serk1-L326F*

Next, we tested whether the *serk1-L326F* mutation interferes with the normal functions of *SERK1.* Prior work has shown that the *serk1-1 serk2-1* double loss-of-function mutant is male sterile due to defective tapetum development and that the *serk1-1 serk2-1 bak1-5* triple mutant is sterile and displays a weak abscission defect (9,27,28). We regenerated the *serk1-1 serk2-1 bak1-5* triple mutant and confirmed the sterility and abscission defective phenotypes [Fig. 2D]. We also created the *serk1-L326F serk2-1 bak1-5* triple mutant and found it displays wildtype fertility and abscises all its floral organs, indicating the *serk1-L326F* mutant allele is a functional co-receptor in tapetum development and abscission [Fig. 2D].

Next, we examined the effect of the *serk1* suppressor mutations in relation to downstream signaling processes. Previously published MAP kinase signaling suppression strategies used in floral abscission research involve inefficient tandem RNAi targeting *MKK4/5* or complicated *MPK3/6* transgenic/mutant combinations (6). Work in stomatal cellular identity specification has shown that MAP kinase signaling regulated by MKK4/5-MPK3/6 can be suppressed in a cell type specific manner by expression of a kinase-inactive version of the MAPKKK *YODA/YDA* gene (29). The identity of the MAPKKK(s) upstream of MKK4/5 during floral abscission is unknown. Nonetheless, we hypothesized that expression of kinase-inactive *YDA* under the control of the *HAE* promoter may block signaling by MKK4/5 during floral abscission.

To test this approach, we created a *HAEpr::YDA-YFP K429R* construct designed to express a YDA-YFP fusion with a mutation of a conserved, catalytic lysine specifically in the abscission zone. We found 33/64 T1 plants transformed with this construct in the Col-0 background displayed abscission defects similar to *hae hsl2* mutants [Fig. 2E; Supp. Table S3]. This phenotype is associated with YFP signal restricted to the abscission zone where HAE is normally expressed [Fig. 2E]. We next tested the ability of this construct to block the suppression effect of *hae-3 hsl2-3 serk1-L326F*. We found that 8/14 T1 *hae-3 hsl2-3 serk1-L326F HAEpr::YDA-YFP K429R* plants had abscission defects similar to *hae hsl2* mutants [Fig. 2E; Supp. Table S3]. These results indicate that the suppression effect acts upstream of MAP kinase signaling, which is consistent with our hypothesis of hypermorphic activation of signaling at the plasma membrane. The results also provide circumstantial evidence that YDA acts downstream of the HAE/HSL2-SERK complex.

### RNA-Sequencing of Col-0, *hae-3 hsl2-3,* and *hae-3 hsl2-3 serk1-L326F*

We next performed an RNA-Sequencing experiment on floral receptacle derived RNA to examine the transcriptome of the *hae-3 hsl2-3 serk1-L326F* mutant in relation to the parental *hae-3 hsl2-3* mutant and the grandparental Col-0. We hypothesized that the *serk1* suppressor would exhibit a reversion of gene expression levels from the *hae-3 hsl2-3* parent toward the Col-0 grandparent [output of differential expression analysis in Supp. Dataset S3].

First, we assessed transcript abundance measurements for the *HAE/HSL2* marker genes *QRT2* and *PGAZAT* [Fig. 3A]. These genes encode polygalacturonases involved in breakdown of pectin in the middle lamella (30). The double *qrt2 pgazat* mutant exhibits a weak abscission delay, and expression of these genes is strongly reduced in the *hae hsl2* mutant (15,30). Thus we consider expression of these genes useful markers for *HAE/HSL2* pathway activity, although it should be noted they are likely only two of many functionally relevant hydrolase genes regulated by *HAE/HSL2.* Consistent with the hypothesis that the *hae-3 hsl2-3 serk1-L326F* mutant is gain of function, the transcript abundance of *QRT2* is increased to an approximately wildtype level [Fig. 3A]. In contrast, *PGAZAT* has a >5-fold increase compared to the wildtype grandparent [Fig. 3A]. These results are consistent with a model where there is activation of a *SERK1* regulated abscission signaling pathway in the *hae-3 hsl2-3 serk1-L326F* mutant.

**Figure 3:**
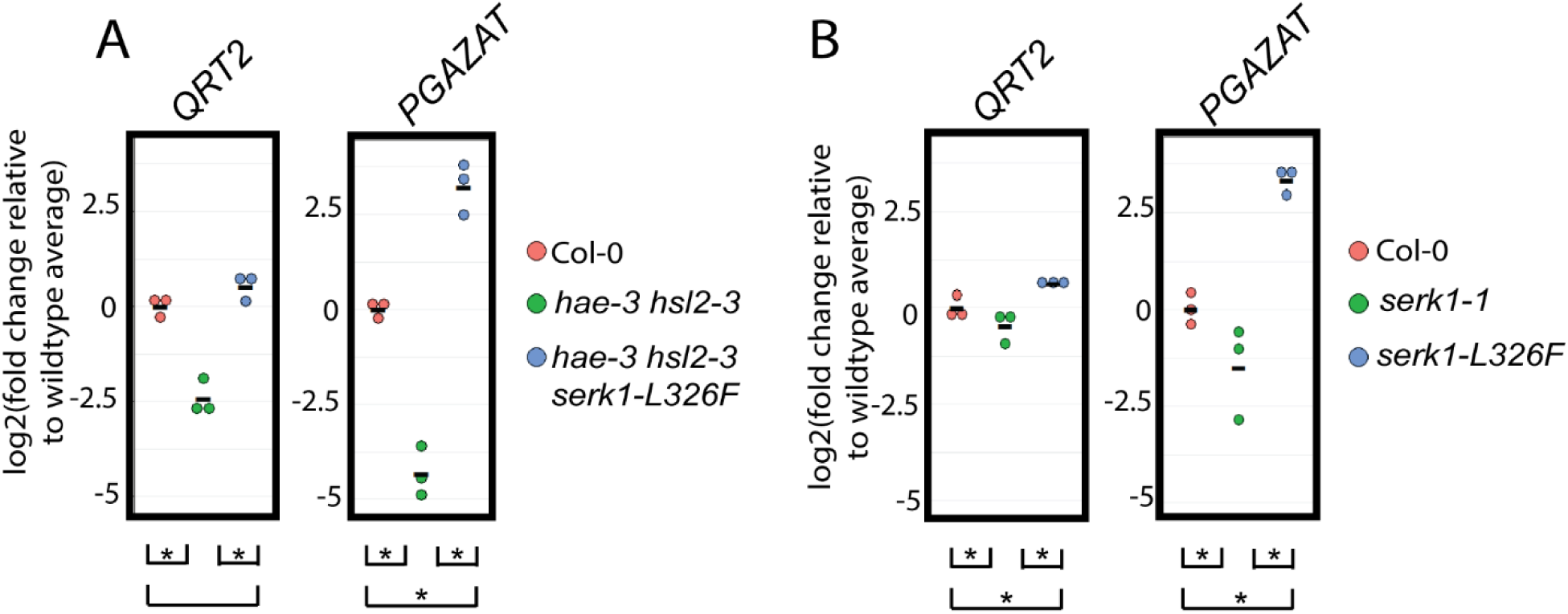
Transcript abundance measurements for abscission hydrolase genes *PGAZAT* and *QRT2*. A) log2(fold change) relative to wildtype average FPKM for abscission hydrolase genes *QRT2* and *PGAZAT*. Asterisk denotes p-value < .05 at FDR of .05. A) log2(fold change) relative to wildtype average FPKM for abscission hydrolase genes *QRT2* and *PGAZAT*. Asterisk denotes p-value < .05 at FDR of .05.

To detect global patterns in gene expression changes in the *hae-3 hsl2-3 serk1-L326F* mutant, we performed Gene Ontology analysis comparing all 3 genotypes [Supp. Table S5]. The most statistically significant findings are terms enriched in the Col-low/*hae-3 hsl2-3 serk1-L326F-*high and *hae-3 hsl2-3*-low/*hae-3 hsl2-3 serk1-L326F-*high comparisons, and include genes associated with terms such as *response to stimulus, response to stress, response to biotic stimulus*, among other terms associated with response to biotic stress [partial list in Table 1]. Overall these results suggest there is a *SERK1* mediated signaling pathway over-activated in *hae-3 hsl2-3 serk1-L326F* regulating both abscission and biotic stress response signaling.

As an additional control, we performed an RNA-Sequencing experiment comparing Col-0 to the single loss of function mutant *serk1-1* and the single putative gain of function mutant *serk1-L326F*. In this experiment, we observed a moderate but statistically significant reduction in expression for *QRT2* and *PGAZAT* in the loss of function *serk1-1* mutant compared to wildtype [Fig. 3B]. The magnitude of this effect was less than that observed in the *hae hsl2* double mutant, consistent with a model where *SERK1* is one of a set of redundant *SERK* genes regulating abscission. In contrast, in the *serk1-L326F* single mutant, we observed a statistically significant increase in both *QRT2* and *PGAZAT* expression compared to wildtype [Fig. 3B]. These results are consistent with a model where *SERK1* positively regulates abscission signaling, and the *serk1-1* and *serk1-L326F* mutations are loss of function and gain of function, respectively.

In addition, in the Col-low/*serk1-L326F* high and *serk1-1-*low/*serk1-L326F* high comparisons, GO analysis identified a strong enrichment in terms such as *response to stimulus, response to stress,* and *response to biotic stimulus*, similar to the *hae-3 hsl2-3 serk1-L326F* mutant [Table 1; Supp. Table S5]. These results are consistent with a model *serk1-L326F* mutant is a gain of function allele broadly activating intracellular signaling. Additionally, we performed quantitative phenotyping of wildtype compared to the single *serk1-1* and *serk1-L326F* mutants [Supp. Fig. S6]. We observed that wildtype abscission occurred at a median floral position between 4 and 5 (floral position 1 is defined as the first flower post-anthesis, with each older flower increasing in position by 1) [Supp. Fig. 6]. We observed the *serk1-L326F* mutant abscising slightly earlier than wildtype at median floral position 4, although this difference was not statistically significant [Supp. Fig. 6]. The single *serk1-1* mutant, in contrast, abscised at a median position of 6, which is highly statistically significantly delayed compared to both wildtype and *serk1-L326F* [Supp. Fig. 6]. These results further support that there is a reduction in signaling in the loss of function *serk1-1* mutant. They also suggest that in *serk1-L326F,* enhanced signaling does not appear to dramatically alter the timing of abscission compared to wildtype. Interestingly, for both *serk1-L236F* single mutant and *hae-3 hsl2-3 serk1-L326F* suppressor mutant, we observed enlarged and disordered abscission zones following abscission by floral position 10, suggesting signaling is not being properly regulated and/or attenuated [Supp. Fig. S6].

The sum of these results indicates that these *SERK1* suppressor mutations are hypermorphic and induce signaling downstream of the plasma membrane through a MAP kinase cascade. This signaling activates broad transcriptional reprogramming, including genes associated with cell wall modification during abscission, as well as biotic stress response signaling.

### Suppression effect does not require SERK1 protein kinase activity

We next investigated potential biochemical mechanisms of *hae hsl2* suppression by the *SERK1* mutations. We hypothesized that the protein kinase domain mutations might over-activate the SERK1 protein kinase domain, causing constitutive phosphorylation of cellular substrates. Contrary to this hypothesis, *in vitro* autophosphorylation analysis did not identify a difference in the autophosphorylation level between wildtype SERK1 and the 3 protein kinase domain mutations with the strongest suppression effect [Fig. 4A]. This result suggests these mutations do not alter intrinsic protein kinase activity of the SERK1 protein.

**Figure 4:**
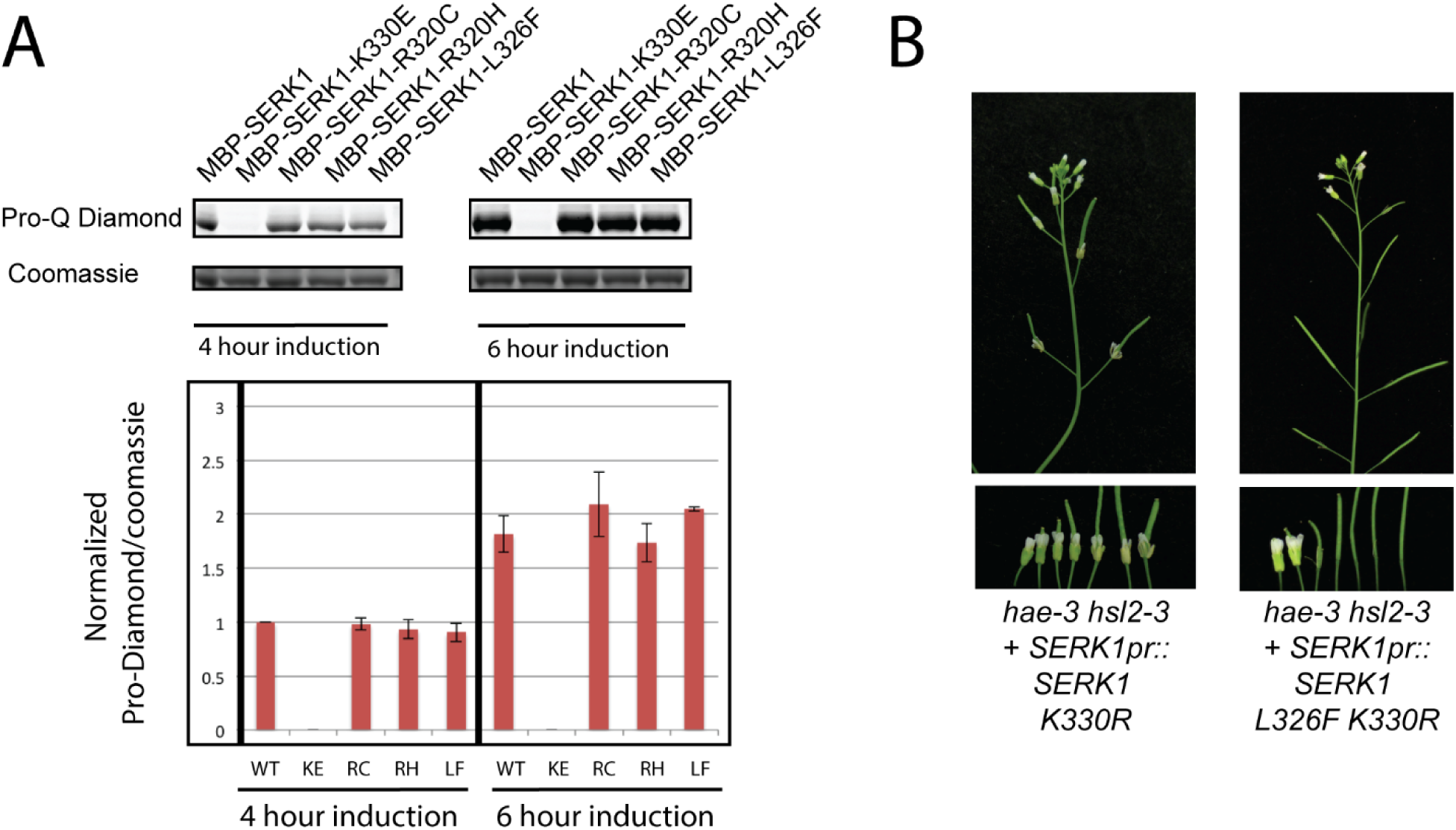
Analysis of kinase activity on *hae-3 hsl2-3* suppression effect. A) *in vitro* autophosphorylation after 4 and 6 hours of induction of recombinant MALTOSE BINDING PROTEIN(MBP)-SERK1 intracellular domain fusion proteins. B) Phenotype of transgenic *hae-3 hsl2-3* transformed with single and double mutant kinase inactive, untagged variants of *SERK1*

We next hypothesized that the *serk1* suppressor mutations alter a negative regulatory interaction *in vivo* that allows SERK1 to phosphorylate cellular substrates constitutively in a manner that does not alter SERK1 protein kinase activity. To test this hypothesis genetically, we used site directed mutagenesis to create a *SERK1pr::SERK1 L326F K330R* transgene. This double mutant lacks an invariant catalytic lysine found in all protein kinases and lacks autophosphorylation activity *in vitro* [Supp. Fig. S7]. We hypothesized that this mutant would be unable to suppress the abscission defective *hae hsl2* phenotype due to abolition of its protein kinase activity. However, in the T1 generation, we found a spectrum of *hae-3 hsl2-3* suppression phenotypes similar to the *SERK1pr::SERK1 L326F* single mutant T1 generation plants (~45% strong or partial suppression in 22 lines) [Fig. 4B; Supp. Table S3]. We discovered this effect also occurs with a version of *SERK1-L326F K330R* tagged with 2xHA [Supp. Fig S7]. These results indicate that phosphorylation of a cellular substrate by the mutant kinase is not the cause of the suppression effect. We speculate that these mutations activate a SERK guarding or monitoring mechanism that senses some biochemical alteration of SERK proteins to transduce a signal downstream, independent of protein kinase activity.

### Evidence that *SOBIR1* transduces signaling downstream of *SERK1* in *hae-3 hsl2-3 serk1-L326F*

We next sought to understand downstream signaling mechanisms in the suppressor mutant. Recent work has shown that overexpression of a *BAK1* transgene lacking the cytosolic kinase domain can activate pathogen response signaling via the RLK *SOBIR1* (31). This result is reminiscent of our observation that signaling by *serk1-L326F* can induce intracellular signaling independent of the protein kinase activity of SERK1. In addition, *SOBIR1* is specifically expressed in abscission zones and has been previously implicated in the regulation of floral abscission (17). Thus we hypothesized *SOBIR1* may function downstream of the suppressor mutations.

To test this model, we crossed the *hae-3 hsl2-3 serk1-L326F* mutant with the exonic *SOBIR1* T-DNA mutant *sobir1-12*/*evr-3*. This mutant allele has previously been reported to suppress the abscission defect of *nev-3* (17,32). We hypothesized that SOBIR1 transduces signaling downstream of SERK1 and thus a *sobir1* mutation would block the effect of the *serk1-L326F* mutation. In the F3 generation, we identified two quadruple homozygous individuals for the four mutations exhibiting strong abscission deficiency [Fig. 5A]. These results suggest *SOBIR1* transduces the signal downstream of the *serk1-L326F* mutation.

**Figure 5:**
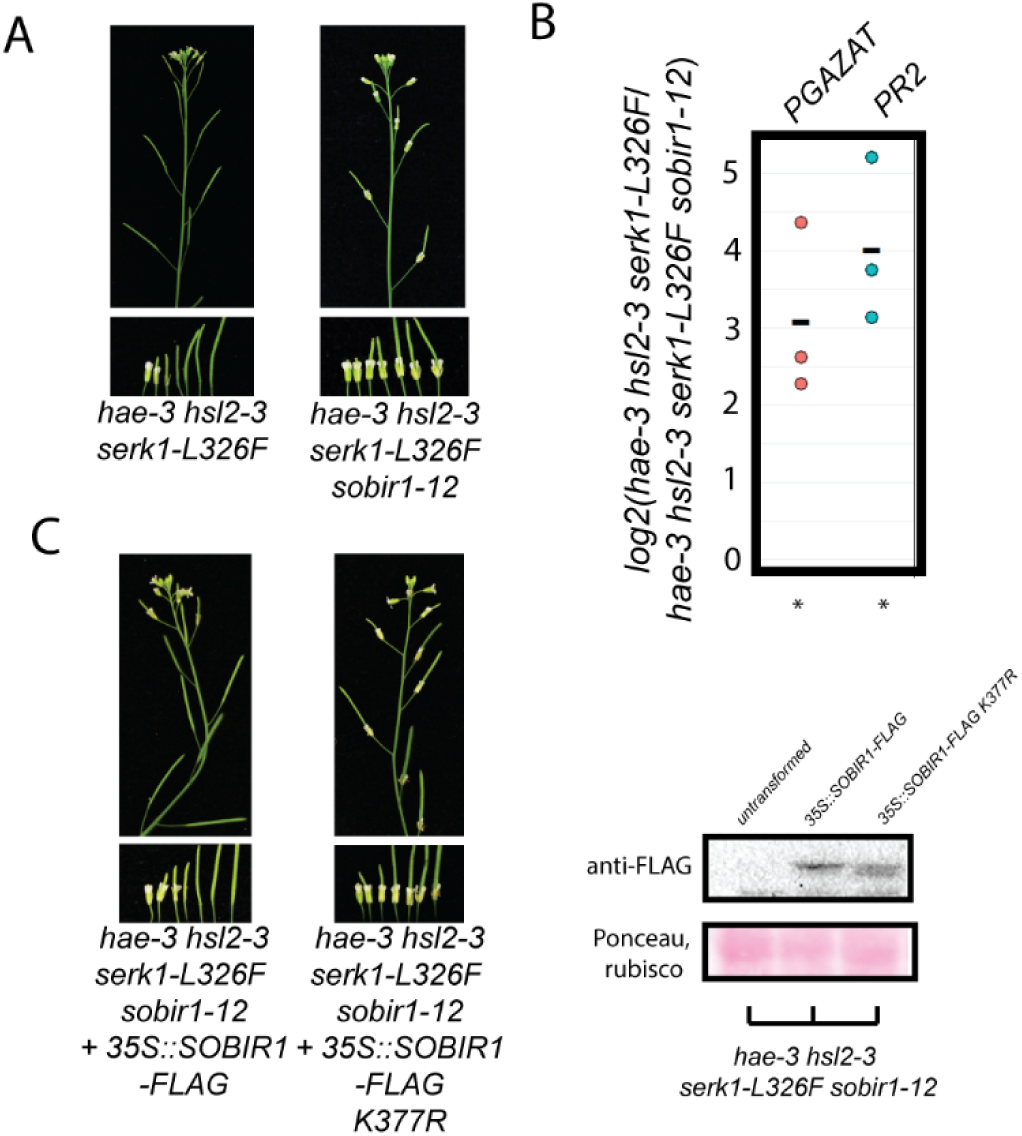
Evidence SOBIR1 functions downstream of *SERK1* during abscission signaling. A) Phenotype of *hae-3 hsl2-3 serk1-L326F* and *hae-3 hsl2-3 serk1-L326F sobir1-12* B) Estimated log2(fold change) of transcript abundance measurements between *hae-3 hsl-3 serk1-L326F* and *hae-3 hsl2-3 serk1-L326F sobir1-12* for abscission hydrolase PGAZAT and pathogen response marker *PR2*. Asterisk denotes p-value < .05. C) Transgenic complementation of the *hae-3 hsl2-3 serk1-L326F sobir1-12* phenotype relies on kinase activity of SOBIR1.

We performed qPCR on the *hae-3 hsl2-3 serk1-L326F sobir1-12* quadruple mutant and compared it with the *hae-3 hsl2-3 serk1-L326F* parent. We utilized a paired design where the difference of the normalized threshold counts for the two genotypes is calculated for each replicate to create a univariate relative gene expression measure. The null hypothesis is that there are no differences between the genotypes and the subtracted threshold counts will be centered around zero. The alternative hypothesis is that the differences will be centered around a non-zero value. We found that the abscission associated hydrolase *PGAZAT* had, on average, an approximately log2(fold change) difference of 3, corresponding to an approximately 8-fold decrease of transcript abundance in the quadruple *hae-3 hsl2-3 serk1-L326F sobir1* mutant [Fig. 5B]. We also tested the pathogen response marker *PR2* and found it exhibited an average log2(fold change) difference of 4, corresponding to an approximately 16-fold decrease in transcript abundance of *PR2* in the quadruple mutant. Overall these results confirm there is a reduction in both abscission and pathogen signaling in the quadruple mutant [Fig. 5B].

To confirm that the quadruple mutant phenotype is due to the mutation in *SOBIR1*, we transformed the quadruple mutant with a transgene expressing either the wildtype *SOBIR1* coding sequence fused to a FLAG tag, or the same transgene with a mutation in the conserved catalytic lysine K377. We observed that the wildtype transgene complemented the abscission deficient phenotype in a majority of T1 lines (58% of 36; Supp. Table S3); whereas, the K377R mutant did not complement any T1 lines examined (0/25; Supp. Table S6) [Fig. 5C]. We screened and found similarly expressing wildtype and K377R lines assayed with an anti-FLAG antibody [Fig. 5C]. Thus, *SOBIR1* can transduce the abscission signal downstream of *serk1-L326F* in a protein kinase activity dependent manner.

### Evidence BIR1 negatively regulates abscission signaling

Next, we sought to understand the mechanism by which the identified *serk1* suppressor mutations might function through *SOBIR1.* The protein kinase BAK1 INTERACTING RLK-1 (BIR1) has been identified as an interactor of SERK proteins (32). In *bir1* mutants, pathogen response signaling is constitutively activated, leading to extreme dwarfism and seedling lethality (32). This effect can be partially suppressed by mutation in *SOBIR1* (32). Thus, BIR1 is thought to negatively regulate signaling by SOBIR1, possibly by acting in a guard complex. Based on these previously known genetic interactions, we hypothesized BIR1 may negatively regulate signaling during abscission and that loss of function of *BIR1* may lead to activation of SOBIR1 in abscission zones in a similar manner to that as *serk1-L326F*.

To test this hypothesis, we sought to combine the *hae hsl2* double mutant with a loss of function *BIR1* mutant. We hypothesized that these plants would over-activate the *SERK1-SOBIR1* abscission pathway and would therefore exhibit restored abscission and enhanced biotic stress response gene expression. *bir1* null mutants are extremely dwarfed and typically die before flowering under standard growth conditions, rendering floral genetic studies difficult. As an alternative, amiRNA targeting *BIR1* has been shown to effectively mimic loss of function mutations in BIR1 (32). Therefore, we created two related *BIR1* amiRNA constructs driven either by the *HAE* promoter alone, or by a tandem *35S::HAEpr*. We anticipated that the *HAEpr* would provide adequate expression levels in the abscission zone, while the tandem *35S::HAEpr* would boost expression in abscission zones in the event that the *HAEpr* proved too weak to be effective.

In the T1, we found that 2/*26 35S::HAEpr::amiRNA-BIR1* plants exhibited partial suppression of the *hae-3 hsl2-3* abscission phenotype, as well as semi-dwarfism with yellowed leaves, while for *HAEpr::amiRNA-BIR1*, 1/16 plants exhibited similar partial abscission and yellow leaves and semi-dwarfism [Fig. 6A; Supp. Table S3]. Thus, loss of *bir1* function appears to suppress the *hae hsl2* abscission defect, phenocopying the gain of function mutation *serk1-L326F.* The leaf phenotype and semi-dwarfism are consistent with weak activation of autoimmunity. To examine these lines, we grew the T2 of one of the *35S::HAEpr* lines and performed qPCR on RNA derived from the floral receptacle on *BIR1* to examine the accumulation of transcript. Across 3 biological replicates, we observed a statistically significant log2(fold change) of .66 for *BIR1* between *hae-3 hsl2-3* and *hae-3 hsl2-3 amiRNA-BIR1*, corresponding to an approximately 40% reduction in *BIR1* transcript accumulation [Fig. 7B]. We also tested transcript abundance of *PGAZAT* and *PR2* and found an approximate log2(fold change) of 5 and 5.4 between *hae-3 hsl2-3* and *hae-3 hsl2-3 amiBIR1* for both genes, respectively, which corresponds to an approximately 32 and 42-fold increase in expression of both genes in the *amiRNA-BIR1* line [Fig. 7B]. These results are consistent with a model where *BIR1* negatively regulates abscission signaling, and that loss of function of *BIR1* leads to high levels of abscission and biotic stress response gene expression, in a manner similar to the *serk1-L326F* mutations.

**Figure 6:**
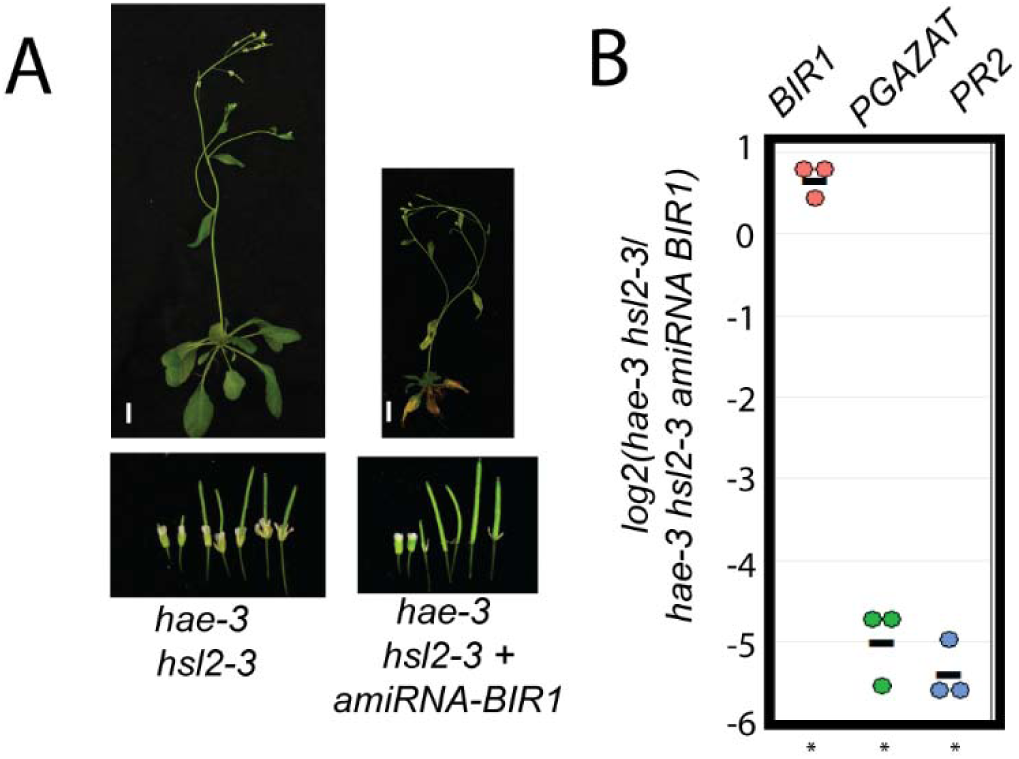
Evidence *BIR1* negatively regulates abscission signaling. A) Phenotype of *hae-3 hsl2-3* and partially suppressed *hae-3 hsl2-3 + HAEpr::amiRNA BIR1*. Scale bar = 1cm B) Estimated log2(fold change) of transcript abundance measurements between *hae-3 hsl2-3* and *hae-3 hsl2-3 amiBIR1* for *BIR1*, abscission hydrolase *PGAZAT*, and pathogen response marker *PR2*. Asterisk denotes p-value < .05.

## Discussion

In this work, we have shown that a particular class of hypermorphic *SERK1* mutations broadly activate intracellular signaling via the RLK *SOBIR1.* The phenotype of these *hae hsl2* suppressor mutations in *SERK1* can be phenocopied by loss of function of the negative regulator of signaling *BIR1.* Prior work has provided strong evidence that the function of BIR1 is to bind to SERK proteins and inhibit activation of multiple pathways, including one regulated by *SOBIR1*. Taken together, these data suggest BIR1 functions in abscission to inhibit overactivation of signaling by SOBIR1. These mutations should provide powerful insight into the activation of SOBIR1, and the regulation of this process by BIR1.

Prior work has shown that *SOBIR1* is specifically expressed in floral abscission zones and that it is globally co-expressed with *HAE* (11,17). This, combined with the genetic data presented in this paper, suggests that *SOBIR1* contributes to wildtype abscission signaling downstream of the SERK proteins. We hypothesize the function of BIR1 during abscission is to moderate SERK mediated activation of SOBIR1 to prevent over-activation of signaling. We further hypothesize that the *serk1* suppressor mutations isolated in this screen interfere with the function of BIR1 by an unknown mechanism. Testing these hypotheses is a clear direction for future research. Notably, the single *sobir1* mutant does not have an abscission defect (17). This implies there are parallel pathways regulating abscission downstream of the HAE/HSL2-SERK complex. An alternative explanation for these results is that, rather than act as a bona fide regulator of abscission signaling, SOBIR1 activation may instead be an undesired byproduct of the activation of the SERK protein signaling complex, and this spurious activation of SOBIR1 may be promoted by the *serk1* suppressor mutations identified in this screen. In this case, the function of BIR1 may simply be to prevent SERK protein mediated activation of SOBIR1 altogether as a way to counteract this inadvertent signal. Given the specific expression pattern of *SOBIR1* in floral abscission zones, this possibility seems unlikely, but is one that cannot be ruled out until additional genetic evidence defining hypothetical parallel pathways including *SOBIR1* come to light.

This work also provides possible insight into the phenotype of the *nevershed* mutant. We have recently shown that the *nev* mutant exhibits widespread transcriptional reprogramming, including extensive induction of biotic stress response gene expression, and that this transcriptional reprogramming is partially reversed in the *nev serk* suppressor line (20). Thus, it appears there may be an aberrant signaling process regulated by *SERK1* which is activated in *nev* that interferes with abscission zone function. This aberrant transcriptional reprogramming is reminiscent of that observed in the *serk1-L326F* mutant. Because loss of function mutations in both *SERK1* and *SOBIR1* can suppress the abscission defect of *nev*, this is evidence that the same pathway activated in the *hae hsl2 serk1* suppressor mutants described in this paper is activated in *nev*. Recent work has shown that the *nev* mutant exhibits over-lignification of the abscission zone (19). This over-lignification may be the result of aberrant signaling in *nev* and could plausibly interfere with normal cell wall modifications required for abscission. Thus, over-lignification represents one possible physical explanation for the phenotype of the *nev* mutant. Why activation of signaling would yield differing phenotypes in *nev* and the *hae hsl2 serk1* suppressor mutants identified in this screen is an important question that will require additional research. It may be that cellular defects of the *nev* mutant render it more susceptible to abscission zone malfunction, or that aberrant signal intensity is higher in *nev,* or a combination of both. Much more work remains to understand *nev* and its suppressors as well as their relationship to the *serk1* mutants identified in this screen, but this work will help guide future investigations in this area.

### Materials and Methods

#### Lines used in this study

*serk1-1* (SALK_044330C), *serk2-1* (SALK_058020C), and *sobir1-12* (SALK_050715) were obtained from the ABRC (17,27,28,32,33). *bak1-5* was kindly provided by Dr. Antje Heese (34). *hae-3 hsl2-3, hae-3 hsl2-3 serk1-L326F, hae-3 hsl2-3 serk1-L326F sobir1-12* have been deposited in the ABRC. The *hae-3 hsl2-3 serk1-L326F* mutant was backcrossed to Col-0 five times. High order mutant combinations were created by cross. All primers used in this study listed in Supp. Table S9.

#### Plant growth conditions

All plants with the exception of the Col-0/*serk1-1/serk1-L326F* RNA-Seq and abscission quantification experiment were grown in a 16 hour light cycle, ~125 μE m-2 s-1, at 22 degrees C. Col-0/*serk1-1/serk1-L326F* RNA-Seq plants were grown in a 16 hour light cycle, ~60 μE m-2 s-1, 23 degrees C. Col-0/*serk1-1/serk1-L326F* abscission quantification plants were grown in a 16 hour light cycle, ~150 μE m-2 s-1, 22 degrees C. All plants were fertilized once at 3 weeks post-germination with 1x strength Miracle-Gro (Scotts Miracle-Gro Company).

#### Phenotyping

Qualitative suppression phenotypes were determined by lightly brushing the main inflorescence of young (<2 weeks post-bolting) plants to remove remnant, abscised floral organs and classifying them according to highest similarity to hae-3 hsl2-3 (non-suppressed), hae-3 hsl2-3 serk1-L326F/+ (partial suppression), and hae-3 hsl2-3 serk1-L326F (strong suppression). Phenotyping of transgenic lines was performed in the T1 generation.

Quantitative phenotyping of Col-0, *serk1-1,* and *serk1-L326F* was performed by utilizing our previously described “petal-puller assay” (35). In this assay, two paint brushes are affixed at an angle to a rigid rod, with spacers present to create a consistent amount of force applied. We lightly dragged the inflorescence of plants at ~2 weeks post-bolting plants, then rotate the plants 90 degrees, and performed the procedure again. The objective of this assay is to remove all remnant floral organs that have undergone abscission but are still loosely attached. Next we counted the number of flowers that had non-abscised floral organs, starting from position 1. The first silique where all floral organs had abscised was recorded. We performed this analysis for 10 plants of each genotype grown in the same flat. We performed a Wilcoxon rank-sum test on the floral positions to test for a shift in location of the median position of abscission.

#### RNA-Sequencing

RNA from between 6 to 15 pooled stage 15 pre-abscission receptacles per replicate was isolated using the Trizol reagent [Life Technologies]. The number of receptacles varied based on the number of stage 15 flowers for each genotype at the time of tissue harvesting. Each receptacle was dissected by taking a 1 mm section of floral tissue comprised of 1/3 mm stem and 2/3 mm receptacle. Libraries were created using the TruSeq mRNA Library Prep Kit [Illumina]. Libraries from each experiment were individually barcoded and run on a single lane of Illumina Sequencing. One replicate of Col-0, *hae-3 hsl2-3,* and *hae-3 hsl2-3 serk1-L326F* was run after the second backcross of the suppressor and *hae-3 hsl2-3* mutants to Col-0 in order to obtain preliminary data, using the Illumina HiSeq 2500. The second and third replicates were performed on the fifth backcrosses and they, along with the Col-0/*serk1-1/serk1-L326F* experiment, were sequenced on an Illumina NextSeq 500.

Reads were mapped to The Arabidopsis Information Resource (TAIR) 10 gene sequences and quantified using TopHat (v. 2.0.9) and Cufflinks (v2.1.1) (36). We utilized default settings for alignments, and performed differential expression analyses using cuffdiff with default settings. Data were analyzed and visualized in R using ggplot2 (36). BAM files have been deposited at Sequence Read Archive under BioProject accession PRJNA430092.

The gene ontology analyses were performed by outputting lists of genes from each comparison found to have a significant difference, with FDR set at .05. These lists were compared using agriGO (37), using Fisher’s exact test, and Yekutieli FDR under dependency, with .05 significance level, against the plant GO database.

#### *hae-3 hsl2-3* suppressor screen

Two suppressor mutant screens were performed on a mutant derived from a cross of *hae-3 hsl2-3* and a previously described *erecta glabra* mutant, both in the Col-0 ecotype (35). *hae-3* contains a missense mutation causing the amino acid residue C222 to be substituted with tyrosine in the extracellular domain, leading to degradation of the mutant protein by an ER associated protein quality control mechanism (21,35). *hsl2-3* contains a missense mutation causing substitution of the amino acid residue G360 with arginine in the HSL2 extracellular domain (21). The molecular defect of *hsl2-3* is presently unknown. We isolated the *hae-3 hsl2-3 er gl* quadruple mutant, which fails to abscise and possesses the characteristic semi-dwarf phenotype of an *erecta* mutant and lacks trichomes. We used this mutant as a background for our screens to control for any wildtype seed contamination that could interfere with suppressor identification. Its short stature makes it convenient for dense planting to screen for floral phenotypes.

We performed an EMS suppressor screen by mutagenizing 50,000 *hae-3 hsl2-3 er gl* seed and growing 50 pools of M1 seeds. Approximately 2,000 seeds from each M1 pool were grown in the M2 to screen for mutant phenotypes. We began initial characterization of four mutants that showed moderate to strong suppression. Two strong suppressors were selected for mapping by crossing to a *hae hsl2* mutant in the Ler ecotype. In the F2, we selected strongly suppressed individuals, pooled the DNA, and sequenced on an Illumina HiSEQ. Reads were aligned with Bowtie2 (version 2.2.6), and SNPs were analyzed with Samtools (version 0.1.19) and the Bar Toronto mutant analysis pipeline to identify a linked region in both mutants on the long arm of chromosome 1 (38–40). Analysis of mutations revealed each mutant had a distinct mutation in the *SERK1* RLK gene. Since *SERK1* is implicated in regulating abscission, it was our highest candidate (26). The other strong mutant and an intermediate mutant were shown, by linkage analysis in the F2 of a cross with Ler *hae hsl2,* to exhibit strong linkage to the marker NGA 111 near the *SERK1* locus [Supp, Fig. 2]. Sanger sequencing the coding regions of *SERK1* in these lines revealed additional missense mutations.

Simultaneously, we performed an activation tagging screen in which 60,000 T1 and 180,000 T2 progeny were screened for suppression of the abscission deficient phenotype of *hae-3 hsl2-3* (41). One line with weak suppression was shown, by TAIL-PCR and by backcross F2 segregation analysis, to have an unlinked T-DNA insertion upstream of At5g09880, a CC1-like splicing factor. However, analysis of the chromatogram of Sanger sequencing of pooled backcross F2 DNA showed a linked SNP in *SERK1* in suppressed individuals, suggesting a spontaneous mutation arose in the *SERK1* gene during creation of this population [Supp. Fig. 2].

#### Molecular cloning

The *SERK1pr::SERK1* construct was created by cloning a ~5 kb fragment of the SERK1 locus including a stop codon into the pE2C entry vector using the NotI site. This was mutagenized by PCR to create *SERK1pr::SERK1-L326F and SERK1pr::SERK1-L326F K330R*. *SERK1pr::SERK1 2xHA* was created by mutagenizing *SERK1pr::SERK1* by PCR based site directed mutagenesis to delete the stop codon and to create an in-frame fusion of *SERK1* with 2xHA. This construct was mutagenized by PCR to create *SERK1pr::SERK1-2xHA L326F*, *SERK1pr::SERK1-2xHA K330R, SERK1pr::SERK1-2xHA L326F K330R*, and *SERK1pr::SERK1-2xHA L326F K330E*. All constructs were combined into the pGWB601 gateway compatible binary vector, transformed into Agrobacterium strain GV3101, transformed into Arabidopsis by floral dip, and selected with Basta (42,43).

*HAEpr::YDA-YFP* kinase inactive was created by cloning the *HAE* promoter into pENTR-TOPO Kinase Inactive-YDA, kindly provided by Dr. Dominique Bergmann (29). This construct was transferred to the binary vector pHGY by gateway recombination, transformed into Arabidopsis by floral dip, and selected with hygromycin on MS agar plates (43,44).

MBP-SERK1 was created by PCR amplifying the intracellular domain of *SERK1* from a cDNA library created from Arabidopsis flowers using Superscript III reverse transcriptase [Thermo Fisher]. This amplicon was cloned into the KpnI site of pMAL-cri and mutagenized by PCR to create pMAL-SERK1-KD-K330E, pMAL-SERK1-KD-L326F, pMAL-SERK1-KD-R330C, pMAL-SERK1-KD-R330H, and pMAL-SERK1-KD-L326F K330R.

The *35Spr::SOBIR1-FLAG* construct was created by PCR amplification of the *SOBIR1* gene from genomic DNA with the addition of a single C-Terminal FLAG tag and stop codon. This fragment was cloned into pENTR/D-TOPO [Thermo Fisher]. PCR based mutagenesis was used to create 35Spr::*SOBIR1-FLAG K377R*. This construct was recombined into the pGWB602 gateway compatible, which contains a 35S driven promoter (42). Plants were transformed by floral dip and selected with Basta.

*HAEpr::amiRNA BIR1* and *35S::HAEpr::amiRNA BIR1* constructs were created by cloning the approximately 2kb upstream of the *HAE* gene into the BamHI/NotI sites of the gateway entry vector pE6c (45). A PacI site was engineered downstream of the *HAEpr*. The *amiRNA BIR1* fragment was created by PCR from primers designed from WMD3 [http://wmd3.weigelworld.org, Ossowski et al, personal communications]. This fragment had a PacI site engineered on the 5’ end. This fragment was PacI digested and cloned into PacI/EcoRV digested pE6c-HAEpr vector. This gateway entry vector was recombined with pGWB601 (*HAEpr::amiRNA BIR1*) or pGWB602 (*35S::HAEpr::amiRNA BIR1*) (42). These constructs were transformed into agrobacterium strain GV3101 and transformed into Arabidopsis by floral dip.

To control for cross contamination, a minimum of one representative T1 individual from each *SERK1* transgenic population was verified by using *hae-3 hsl2-3* dCAPs markers [Supp. Table 4] and by sequencing the *SERK1* transgene across the mutation site(s) by analysis of a PCR product using *SERK1* F primer 5’-TGGAACAACTGTTAATGAAAATCAA-3’ and pE2c R 5’-AGTCGGGCACGTCGTAGG-3’.

A PCR product from a representative T1 individual from each *HAEpr::YDA-YFP K429R* population was subjected to Sanger sequencing using a HAE promoter specific primer 5’-AGGCAGAGTGCTTGTGGAGACG-3’ and a YDA specific primer 5’-CAGGTGCCATCCAATATGGGCTC-3’. The *hae-3 hsl2-3 serk1-L326F* T1 individual also was subjected to genotyping with *serk1-L326F* dCAPS primers [Supp. Table 4].

The strongly expressing *hae-3 hsl2-3 serk1-L326F sobir1-12* transgenic individuals were validated by genotyping with *hae-3, hsl2-3* and *serk1-L326F* dCAPS markers as well as *SOBIR1* wildtype and T-DNA specific primers.

A single T2 *hae-3 hsl2-3 + 35S::HAEpr::amiRNA BIR1* plant was validated by *hae-3* and *hsl2-3* dCAPS markers and by PCR amplifying and Sanger sequencing across the amiRNA sequence using *HAEpr* primer 5’-TTCACATGGATGTATACTATTGCCTCCT-3’ and amiRNA primer B 5’-GCGGATAACAATTTCACACAGGAAACAG-3’ to ensure it correctly aligned to the output generated by WMD3.

All constructs were created using PFU Ultra II High-Fidelity polymerase and Sanger sequenced verified across the entire coding region [Agilent]. All primers are listed in Supplemental Table 4.

#### Western blotting

Three whole flowers, starting with a stage 15 flower and proceeding with the next two oldest flowers, were frozen and ground in microcentrifuge tubes, resuspended in 30 µl of SDS sample buffer, and boiled for ~3 min. 10 µl of each sample was run on an 8% acrylamide gel, blotted to a nitrocellulose membrane, stained with Ponceau-S, and imaged. Blots were blocked with 4% BSA in phosphate-buffered saline with 0.1% Tween-20 (PBS-T) for 1 h, probed with anti-HA-HRP or anti-FLAG antibody at 1:1000 dilution for 1 hour at room temperature or overnight at 4 degrees C, respectively, then rinsed with PBS-T four times for 5 min each [anti-HA-HRP antibody Roche clone 3F10, anti-FLAG antibody Sigma M2]. HA blots were directly imaged. FLAG blots were incubated with anti-mouse–horseradish peroxidase (HRP) (1:2500 dilution in 1% BSA in PBS-T, Cell Signaling Technologies, Danvers, MA, USA) for 1 h at room temperature, rinsed with PBS-T four times for 5 min each. Blots were imaged by incubation with a chemiluminescent substrate (SuperSignal West Pico, Life Technologies, Carlsbad, CA, USA) and imaged using a Bio-Rad ChemiDoc.

#### qPCR

RNA from five stage 15 receptacles was isolated using the Trizol reagent, from which cDNA was synthesized following DNAse I treatment. Three or five biological replicates per reaction were performed. We used ABsolute SYBR Green Master Mix (2X) [Thermo Fisher] and a Bio-Rad CFX96 thermal cycler to estimate threshold counts during the log-linear phase of amplification. Data analysis was performed in Microsoft Excel. A paired t-test was performed on the raw delta-delta-Ct between the gene of interest and previously published reference gene(s) for each replicate (46,47). We utilized reference gene AT5G25760 for normalization for *hae-3 hsl2-3 serk1-L326F sobir1-12* experiment and the geometric mean of AT3G01150 and AT2G28390 for *BIR1* experiments (48). An 8-fold serial dilution was used to calculate PCR primer efficiency and determine estimated fold change levels.

#### *in vitro* autophosphorylation

*In vitro* autophosphorylation assays were carried out exactly as described in (49). In brief, expression of MBP-SERK1 was induced by adding isopropyl-β-d-thiogalactopyranoside (IPTG) to rapidly growing *E. coli* to a final concentration of 0.1 mM. These cultures were then allowed to grow for 4 or 6 hours, after which 100 µl of liquid culture was spun down, resuspended in 100 µl of 1x sodium dodecyl sulfate (SDS) sample buffer, and boiled for 3 min. 10 µl of the whole cell lysate was then run on an 8% acrylamide gel after which the gel was fixed by incubation in 50% ethanol/10% acetic acid overnight. The gel was rehydrated by soaking in DdH20 two times for 30 min and then immersed in one-third strength Pro-Q Diamond dye for 2 h in the dark on a rotating platform [Thermo Fisher]. The gel was destained in 20% acetonitrile, 50 mM sodium acetate (pH 4.0) four times for 30 min each time. Then, it was soaked in DdH20 twice for 10 min before imaging on a Bio-Rad ChemiDoc using the default Pro-Q Diamond settings. Finally, the gel was stained with Coomassie and imaged using a Bio-Rad GelDoc. Band quantity estimates were performed using the built in Bio-Rad software.

## Acknowledgements

We would like to greatly thank the skilled staff at the MU DNA Core Facility for Illumina Sequencing services. We also would like to thank Melody Kroll for thoughtful review of the manuscript. We thank Dr. Tsuyoshi Nakagawa (Shimane University) for providing the pGWB601/pGWB602 binary vectors that contains the bar gene, which was identified by Meiji Seika Kaisha, Ltd. The Gateway entry vector pE2C was obtained from Addgene.

## Author contributions

Isaiah Taylor designed, performed, and analyzed the results of experiments and wrote the paper. John Baer performed the *in vitro* autophosphorylation assays. Ryan Calcutt performed genetic analyses. John C. Walker oversaw the work and edited the paper. All authors edited and approved the final version of the paper.

## References

1. Patterson SE. Cutting Loose. Abscission and Dehiscence in Arabidopsis. Plant Physiol. 2001 Jun 1;126(2):494–500.

2. Liljegren SJ, Leslie ME, Darnielle L, Lewis MW, Taylor SM, Luo R, et al. Regulation of membrane trafficking and organ separation by the NEVERSHED ARF-GAP protein. Development. 2009;136(11):1909–1918.

3. Patharkar OR, Walker JC. Core Mechanisms Regulating Developmentally Timed and Environmentally Triggered Abscission. Plant Physiol. 2016 Sep;172(1):510–20.

4. Patharkar OR, Gassmann W, Walker JC. Leaf shedding as an anti-bacterial defense in Arabidopsis cauline leaves. PLoS Genet. 2017;13(12):e1007132.

5. Jinn TL, Stone JM, Walker JC. HAESA, an Arabidopsis leucine-rich repeat receptor kinase, controls floral organ abscission. Genes Dev. 2000 Jan 1;14(1):108–17.

6. Cho SK, Larue CT, Chevalier D, Wang H, Jinn T-L, Zhang S, et al. Regulation of floral organ abscission in Arabidopsis thaliana. Proc Natl Acad Sci U S A. 2008 Oct 7;105(40):15629–34.

7. Stenvik G-E, Tandstad NM, Guo Y, Shi C-L, Kristiansen W, Holmgren A, et al. The EPIP peptide of INFLORESCENCE DEFICIENT IN ABSCISSION is sufficient to induce abscission in arabidopsis through the receptor-like kinases HAESA and HAESA-LIKE2. Plant Cell. 2008 Jul;20(7):1805–17.

8. Schardon K, Hohl M, Graff L, Pfannstiel J, Schulze W, Stintzi A, et al. Precursor processing for plant peptide hormone maturation by subtilisin-like serine proteinases. Science. 2016;354(6319):1594–1597.

9. Meng X, Zhou J, Tang J, Li B, de Oliveira MV, Chai J, et al. Ligand-induced receptor-like kinase complex regulates floral organ abscission in Arabidopsis. Cell Rep. 2016;14(6):1330–1338.

10. Santiago J, Brandt B, Wildhagen M, Hohmann U, Hothorn LA, Butenko MA, et al. Mechanistic insight into a peptide hormone signaling complex mediating floral organ abscission. Elife. 2016;5:e15075.

11. Patharkar OR, Walker JC. Floral organ abscission is regulated by a positive feedback loop. Proc Natl Acad Sci. 2015;201423595.

12. Fernandez DE, Heck GR, Perry SE, Patterson SE, Bleecker AB, Fang S-C. The embryo MADS domain factor AGL15 acts postembryonically: inhibition of perianth senescence and abscission via constitutive expression. Plant Cell. 2000;12(2):183–197.

13. Adamczyk BJ, Lehti-Shiu MD, Fernandez DE. The MADS domain factors AGL15 and AGL18 act redundantly as repressors of the floral transition in Arabidopsis. Plant J. 2007;50(6):1007–1019.

14. Chen M-K, Hsu W-H, Lee P-F, Thiruvengadam M, Chen H-I, Yang C-H. The MADS box gene, FOREVER YOUNG FLOWER, acts as a repressor controlling floral organ senescence and abscission in Arabidopsis. Plant J. 2011;68(1):168–185.

15. Niederhuth CE, Patharkar OR, Walker JC. Transcriptional profiling of the Arabidopsis abscission mutant hae hsl2 by RNA-Seq. BMC Genomics. 2013;14:37.

16. Butenko MA, Patterson SE, Grini PE, Stenvik G-E, Amundsen SS, Mandal A, et al. INFLORESCENCE DEFICIENT IN ABSCISSION controls floral organ abscission in Arabidopsis and identifies a novel family of putative ligands in plants. Plant Cell Online. 2003;15(10):2296–2307.

17. Leslie ME, Lewis MW, Youn J-Y, Daniels MJ, Liljegren SJ. The EVERSHED receptor-like kinase modulates floral organ shedding in Arabidopsis. Dev Camb Engl. 2010 Feb;137(3):467–76.

18. Burr CA, Leslie ME, Orlowski SK, Chen I, Wright CE, Daniels MJ, et al. CAST AWAY, a membrane-associated receptor-like kinase, inhibits organ abscission in Arabidopsis. Plant Physiol. 2011 Aug;156(4):1837–50.

19. Lee Y, Yoon TH, Lee J, Jeon SY, Lee JH, Lee MK, et al. A lignin molecular brace controls precision processing of cell walls critical for surface integrity in Arabidopsis. Cell. 2018;173(6):1468–1480.

20. Taylor I, Walker JC. Transcriptomic evidence for distinct mechanisms underlying abscission deficiency in the Arabidopsis mutants haesa/haesa-like 2 and nevershed. BMC Res Notes. 2018 Oct 23;11(1):754.

21. Niederhuth CE, Cho SK, Seitz K, Walker JC. Letting Go is Never Easy: Abscission and Receptor-Like Protein Kinases. J Integr Plant Biol. 2013;55(12):1251–1263.

22. Yan L, Ma Y, Liu D, Wei X, Sun Y, Chen X, et al. Structural basis for the impact of phosphorylation on the activation of plant receptor-like kinase BAK 1. Cell Res. 2012;22(8):1304.

23. Santiago J, Henzler C, Hothorn M. Molecular mechanism for plant steroid receptor activation by somatic embryogenesis co-receptor kinases. Science. 2013;341(6148):889–892.

24. Jaillais Y, Belkhadir Y, Balsemão-Pires E, Dangl JL, Chory J. Extracellular leucine-rich repeats as a platform for receptor/coreceptor complex formation. Proc Natl Acad Sci. 2011;108(20):8503–8507.

25. Hohmann U, Nicolet J, Moretti A, Hothorn LA, Hothorn M. The SERK3 elongated allele defines a role for BIR ectodomains in brassinosteroid signalling. Nat Plants. 2018;4(6):345.

26. Lewis MW, Leslie ME, Fulcher EH, Darnielle L, Healy PN, Youn J-Y, et al. The SERK1 receptor-like kinase regulates organ separation in Arabidopsis flowers. Plant J Cell Mol Biol. 2010 Jun 1;62(5):817–28.

27. Colcombet J, Boisson-Dernier A, Ros-Palau R, Vera CE, Schroeder JI. Arabidopsis SOMATIC EMBRYOGENESIS RECEPTOR KINASES1 and 2 are essential for tapetum development and microspore maturation. Plant Cell Online. 2005;17(12):3350–3361.

28. Albrecht C, Russinova E, Hecht V, Baaijens E, de Vries S. The Arabidopsis thaliana SOMATIC EMBRYOGENESIS RECEPTOR-LIKE KINASES1 and 2 control male sporogenesis. Plant Cell Online. 2005;17(12):3337–3349.

29. Lampard GR, Lukowitz W, Ellis BE, Bergmann DC. Novel and Expanded Roles for MAPK Signaling in Arabidopsis Stomatal Cell Fate Revealed by Cell Type–Specific Manipulations. Plant Cell Online. 2009;21(11):3506–3517.

30. Ogawa M, Kay P, Wilson S, Swain SM. ARABIDOPSIS DEHISCENCE ZONE POLYGALACTURONASE1 (ADPG1), ADPG2, and QUARTET2 are polygalacturonases required for cell separation during reproductive development in Arabidopsis. Plant Cell. 2009;21(1):216–233.

31. Domínguez-Ferreras A, Kiss-Papp M, Jehle AK, Felix G, Chinchilla D. An overdose of the Arabidopsis coreceptor BRASSINOSTEROID INSENSITIVE1-ASSOCIATED RECEPTOR KINASE1 or its ectodomain causes autoimmunity in a SUPPRESSOR OF BIR1-1-dependent manner. Plant Physiol. 2015;168(3):1106–1121.

32. Gao M, Wang X, Wang D, Xu F, Ding X, Zhang Z, et al. Regulation of cell death and innate immunity by two receptor-like kinases in Arabidopsis. Cell Host Microbe. 2009 Jul 23;6(1):34–44.

33. Alonso JM, Stepanova AN, Leisse TJ, Kim CJ, Chen H, Shinn P, et al. Genome-wide insertional mutagenesis of Arabidopsis thaliana. Science. 2003;301(5633):653–657.

34. Schwessinger B, Roux M, Kadota Y, Ntoukakis V, Sklenar J, Jones A, et al. Phosphorylation-dependent differential regulation of plant growth, cell death, and innate immunity by the regulatory receptor-like kinase BAK1. PLoS Genet. 2011;7(4):e1002046.

35. Baer J, Taylor I, Walker JC. Disrupting ER-associated protein degradation suppresses the abscission defect of a weak hae hsl2 mutant in Arabidopsis. J Exp Bot. 2016;67(18):5473–5484.

36. Trapnell C, Roberts A, Goff L, Pertea G, Kim D, Kelley DR, et al. Differential gene and transcript expression analysis of RNA-seq experiments with TopHat and Cufflinks. Nat Protoc. 2012;7(3):562.

37. Du Z, Zhou X, Ling Y, Zhang Z, Su Z. agriGO: a GO analysis toolkit for the agricultural community. Nucleic Acids Res. 2010;38(suppl_2):W64–W70.

38. Langmead B, Salzberg SL. Fast gapped-read alignment with Bowtie 2. Nat Methods. 2012;9(4):357–359.

39. Li H, Handsaker B, Wysoker A, Fennell T, Ruan J, Homer N, et al. The sequence alignment/map format and SAMtools. Bioinformatics. 2009;25(16):2078–2079.

40. Austin RS, Vidaurre D, Stamatiou G, Breit R, Provart NJ, Bonetta D, et al. Next-generation mapping of Arabidopsis genes. Plant J. 2011;67(4):715–725.

41. Weigel D, Ahn JH, Blázquez MA, Borevitz JO, Christensen SK, Fankhauser C, et al. Activation Tagging in Arabidopsis. Plant Physiol. 2000 Apr 1;122(4):1003–14.

42. Nakamura S, Mano S, Tanaka Y, Ohnishi M, Nakamori C, Araki M, et al. Gateway Binary Vectors with the Bialaphos Resistance Gene, *bar*, as a Selection Marker for Plant Transformation. Biosci Biotechnol Biochem. 2010;74(6):1315–9.

43. Clough SJ, Bent AF. Floral dip: a simplified method forAgrobacterium-mediated transformation ofArabidopsis thaliana. Plant J. 1998;16(6):735–743.

44. Kubo M, Udagawa M, Nishikubo N, Horiguchi G, Yamaguchi M, Ito J, et al. Transcription switches for protoxylem and metaxylem vessel formation. Genes Dev. 2005 Aug 15;19(16):1855–60.

45. Dubin MJ, Bowler C, Benvenuto G. A modified Gateway cloning strategy for overexpressing tagged proteins in plants. Plant Methods. 2008;4(3):1–11.

46. Yuan JS, Reed A, Chen F, Stewart CN. Statistical analysis of real-time PCR data. BMC Bioinformatics. 2006;7(1):85.

47. Dussault A-A, Pouliot M. Rapid and simple comparison of messenger RNA levels using real-time PCR. Biol Proced Online. 2006;8(1):1–10.

48. Czechowski T, Stitt M, Altmann T, Udvardi MK, Scheible W-R. Genome-wide identification and testing of superior reference genes for transcript normalization in Arabidopsis. Plant Physiol. 2005;139(1):5–17.

49. Taylor I, Seitz K, Bennewitz S, Walker JC. A simple in vitro method to measure autophosphorylation of protein kinases. Plant Methods. 2013;9(1):22.

